# Safety first: input screening for protein design tools

**DOI:** 10.64898/2026.08.04.740855

**Authors:** Phil Palmer, Nikki Teran, Nicole Wheeler, Jaime Yassif

## Abstract

As biological AI models become more powerful, practical biosecurity approaches are needed to support beneficial applications while reducing misuse risks. Sequence-similarity-based screening approaches are no longer adequate to safeguard biological AI models because these models can design molecules with novel sequences and structures. Therefore, a screening approach that takes function into account is needed. To address this need, we propose a new screening method for AI-enabled protein binder design tools. Our framework screens protein binding targets, with a focus on the human proteome, as opposed to the binder molecule itself. We constructed a database of 14,541 potentially harmful proteoform targets from the human proteome (7.1% of all human protein proteoforms) classified by biosecurity risk level. To discern structural and functional features, we evaluated constructs with an embedding-based screening method using the ESM-C protein language model. ESM-C achieved high accuracy for detecting variants of known targets (F1 scores >97%), with performance similar to BLASTP. However, ESM-C proved to be more effective at capturing functional relationships, distinguishing benign mutations from damaging ones where BLASTP did not.

To characterize how screening would affect bioscience research, we measured flagging rates across diverse protein datasets. Flagging rates were significant for mammalian proteins weighted by publication frequency (23% for human, 20% for mouse), and rates for organisms distantly related to humans were minimal (<1.1% for bacteria, fungi, plants, and viruses). Among commercially relevant targets, 63% of antibody patent targets were classified as dual-use, reflecting that therapeutically important proteins often perform critical biological functions. To identify and flag risky user requests from protein binder design tools without placing an undue burden on scientific research and innovation, it will be essential to deploy this screening approach in a way that addresses the overlap our analysis showed between targets of concern and therapeutic targets–possibly in concert with tiered trusted access frameworks. This new method provides a foundation for proportionate safeguards for biological AI models that reduce misuse risks while preserving their benefits for legitimate research and demonstrates a concrete proof of principle that can be generalized to other protein design tools and biological AI models.

## 1 Introduction

### Biological AI models

Recent advances have produced biological AI models (BAIMs) with transformative potential for human health and the bioeconomy. AlphaFold^1^ and ESMFold^2^ predict protein structures from sequences, accelerating drug discovery and enabling rational protein engineering. Protein language models such as ESM^3^ learn rich numerical representations of proteins that capture functional and evolutionary relationships, enabling applications from variant effect prediction to protein design. Protein binder design tools generate novel proteins that bind specified targets, with applications in therapeutics, diagnostics, and research.^4–10^

These same capabilities also create biosecurity risks: tools designed to create therapeutic proteins targeting the human proteome could equally design novel pathogen or toxin proteins that disrupt critical biological functions. According to Epoch AI, only 3.2% of BAIMs have any safety measures in place,^11^ although the majority of current BAIMs are low risk. The burden of implementing safeguards currently falls to individual tool developers, resulting in protections that are inconsistent and mostly absent.

### Protein binder design tools

Protein binder design tools generate novel proteins that bind to a user-specified target, potentially disrupting its function. Most accept the target protein structure as input (AlphaProteo,^4^ BindCraft,^5^ BoltzGen,^6^ Chai-2,^7^ RFantibody,^9^ RFdiffusion^10^), though some, such as EvoBind2,^8^ require only the target sequence.

We focus on protein binder design tools because they represent a well-defined category within the broader class of BAIMs, making them a natural starting point for developing a screening framework. Future work can extend this approach to other tool categories.

### Addressing gaps in biosecurity screening

This tool is designed to address a critical gap in biosecurity screening. While there has been extensive investment in DNA synthesis screening, which operates at the interface between digital designs and physical biological systems, no tools currently exist to screen user requests to protein design tools.

Existing biosecurity screening also relies primarily on sequence alignment methods such as BLAST,^12,13^ which can only detect sequences similar to those already in a database. Recent work has demonstrated that AI-designed proteins can have low sequence similarity to natural proteins while retaining similar function,^14,15^ highlighting the need for structural and functional screening approaches. Prior to this work, no database of concerning protein targets existed, and no screening framework had been proposed for protein design tools.

### Contributions

We propose a framework for screening protein binder design tools that focuses on inputs (user design requests) rather than outputs (designed sequences) (Figure 1). Our input screening approach identifies targets of concern (TOC), rather than screening sequences of concern (SOC), the standard approach for DNA synthesis screening. As an example of this approach, SARS-CoV-2 enters human cells by binding the ACE2 receptor via its Spike protein; input screening would flag a request to design a binder targeting ACE2 rather than attempting to screen the designed sequence after generation.

**Figure 1:**
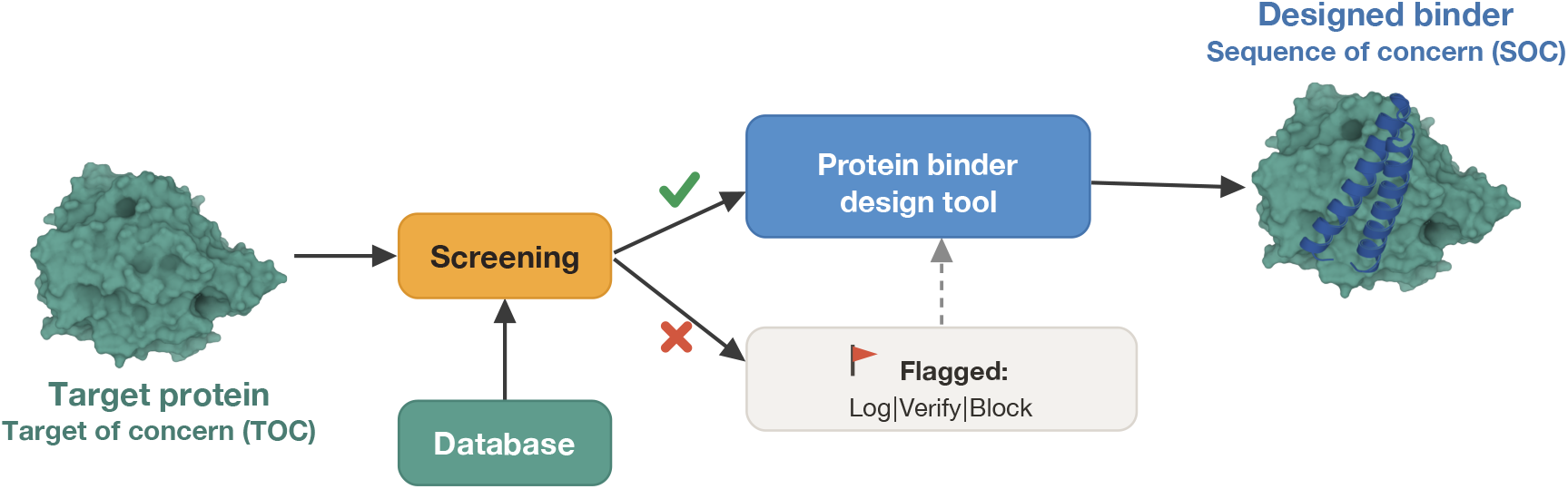
Screening workflow for protein design tools. Input target proteins are screened against a database of targets of concern before design proceeds. Flagged requests can be blocked or logged; unflagged requests proceed to binder design.

Input screening has several advantages. Importantly, screening occurs before computation, avoiding wasted resources on requests that are subsequently blocked. Additionally, the parameter space of possible binding targets is constrained to existing proteins rather than the vast space of designed sequences. For a protein of even modest length (75 residues), the number of possible sequences exceeds the estimated number of atoms in the observable universe,^15^ the minority being either known-benign or known-hazardous sequences and the majority being novel and indeterminate. This makes comprehensive database coverage feasible only for input screening.

We systematically analyze the entire human proteome (both reviewed and unreviewed proteins) to construct a database of 14,541 targets of concern (7.1% of the proteoforms from UniProt) for testing our methods. This paper focuses primarily on the target-of-concern and embedding-space-based methods, and less on the database creation and optimization. In principle these methods can be applied to any database with targets of concern. For the purposes of the following analysis, we classify the targets of concern by risk level. Targets with high harm potential with limited therapeutic benefit were classified as red, while dual-use targets with high harm potential that also are known therapeutic targets were classified as yellow, given their beneficial and potentially dangerous applications. We also classified targets as yellow if they had moderate harm potential irrespective of therapeutic benefit. This database can, and should, be optimized and updated as biomedical knowledge evolves.

We develop an embedding-based screening method using protein language models and compare performance to BLASTP to evaluate whether embedding-based approaches offer advantages over established sequence alignment methods. We investigate robustness to sequence variation and characterize flagging rates across diverse research contexts, with the goal of improving biosecurity while minimizing disruption to beneficial applications.

## 2 Results

### 2.1 Overview of the screening approach

The screening method uses ESM-C, a protein language model that generates numerical representations (referred to as embeddings) of proteins.^16^ Functionally similar proteins have similar embeddings. For each incoming design request, we compute the embedding of the target protein and compare it to pre-computed embeddings for all 14,541 targets of concern in the human proteome database (see Methods for database construction). If the maximum cosine similarity, which measures relatedness between two protein embeddings, exceeds the threshold, described in section 2.2, the request is flagged.

To visualize this approach, we projected ESM-C embeddings into two dimensions using uniform manifold approximation and projection (UMAP) (Figure 2A). Targets of concern (shown in red) and therapeutic targets (shown in blue) show substantial overlap in the UMAP projection, which is consistent with the dual-use nature of many biologically important proteins. Despite this overlap in the wider UMAP projection, the screening threshold distinguishes most targets: the zoomed region (Figure 2B) shows that most therapeutic targets, even in this projection space, fall outside the flagging threshold, though some are flagged due to proximity to targets of concern. All analysis is performed in the higher-dimension embedding space; the UMAP projections only serve as an approximate visualization here.

**Figure 2:**
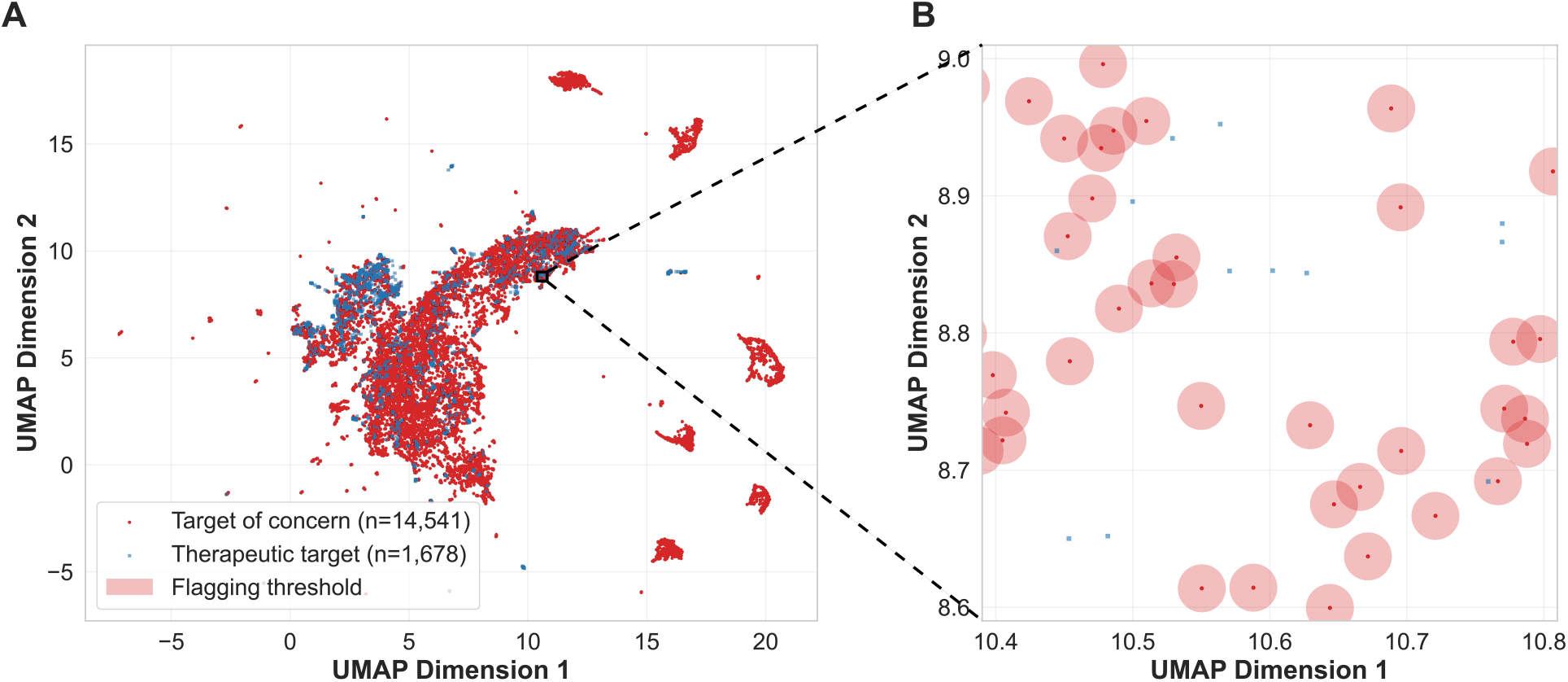
Embedding-based screening in protein embedding space. (A) UMAP projection of ESM-C embeddings for targets of concern (red, n=14,541) and clinically validated therapeutic targets (blue, n=1,678). (B) Zoomed region illustrating the screening threshold. Semi-transparent circles approximate the cosine similarity flagging threshold (0.9903) around each target of concern. Circle radii were estimated from the median UMAP distance between threshold-adjacent therapeutic proteins and their nearest target of concern. Note: The threshold boundaries shown are approximations as UMAP reduces 1,152 dimensions to two and distorts distances non-uniformly; screening uses cosine similarity in the full-dimensional space.

### 2.2 Threshold optimization and baseline performance

To calibrate the screening system, we optimized similarity thresholds for ESM-C using a dataset designed to test the core screening task: distinguishing variants of targets of concern from therapeutic targets. We also compared performance of ESM-C to BLAST, specifically BLASTP which compares protein sequences, to test whether, under a particular set of conditions, its performance is comparable to this sequence-screening-based approach.

The test set consisted of two classes: benign triple-SNV (single nucleotide variant) variants of targets of concern, which should be flagged as they represent minor natural variation of concerning proteins; and clinically validated therapeutic targets with no direct overlap with red or yellow database categories, which should pass screening. We optimized thresholds on a calibration set to maximize F1 score (Figure 3A–B). For the ESM-C embedding, the optimal threshold of 0.9903 maximizes F1 score at 96.9%. This threshold is used in the embedding-based screening approach described above. However, this triple-SNV threshold is more stringent than may be relevant for real-world, biosecurity-relevant screening where obfuscation may be intentional.

**Figure 3:**
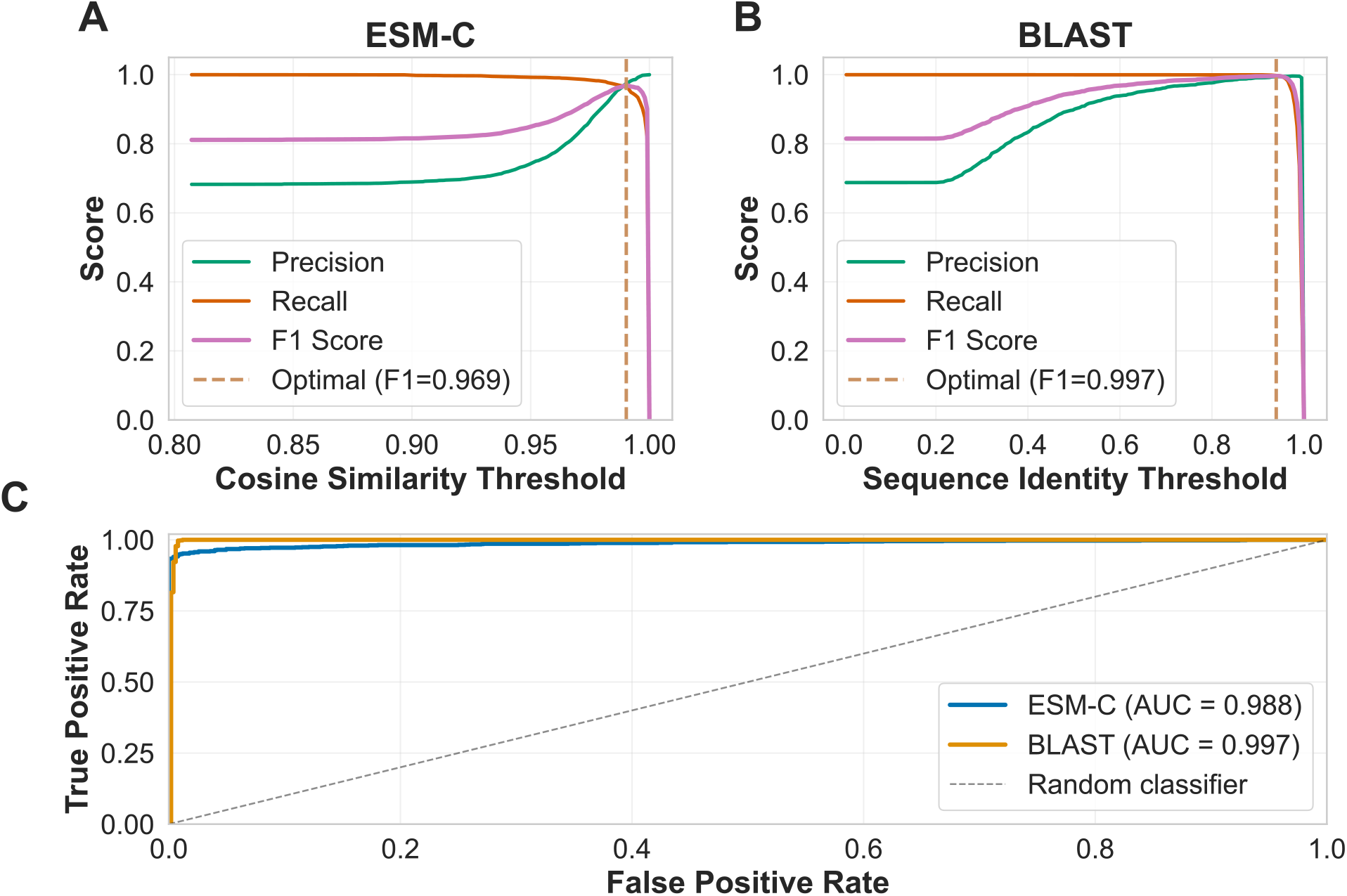
Threshold optimization for embedding-based and alignment-based screening. (A) Precision (green), recall (orange), and F1 score (pink) as a function of cosine similarity threshold for ESM-C screening on the calibration dataset. The optimal threshold (0.9903, brown dashed line) maximizes F1 score at 96.9%. (B) Equivalent analysis for BLASTP screening using protein sequence identity. The optimal threshold (0.9397, brown dashed line) maximizes F1 score at 99.7%. (C) Receiver operating characteristic curves on the test dataset comparing ESM-C (AUC = 0.988) and BLASTP (AUC = 0.997). The evaluation dataset consisted of triple-SNV variants of targets of concern (calibration n=2,520, test n=1,081) and clinically validated therapeutic targets with no database overlap (calibration n=1,174, test n=504). Values are for a single 70/30 calibration/test split. Performance is stable across 100 resampled splits (F1 96.9 ± 0.3% ESM-C, 99.6 ± 0.1% BLASTP; AUC 98.5 ± 0.2% and 99.4 ± 0.3%), with the ESM-C optimal threshold varying by only ±0.0004. The small F1 difference is consistent across splits (Nadeau–Bengio corrected resampled t-test, p < 0.001).

We then evaluated final performance on a held-out test set. Both ESM-C and BLASTP methods achieved high accuracy on this test set (Table 1, Figure 3C).

**Table 1.** Screening performance at optimal thresholds on the test set.

| Method | Threshold | F1 score | Precision | Recall | AUC |
| --- | --- | --- | --- | --- | --- |
| ESM-C (cosine similarity) | 0.9903 | 97.2% | 98.0% | 96.4% | 98.8% |
| BLASTP (protein sequence identity) | 0.9397 | 99.7% | 99.7% | 99.7% | 99.7% |

**Table 2.** *Risk tiers*. Reviewed status is taken from UniProt.

| Tier | Count | % of proteome | % of reviewed |
| --- | --- | --- | --- |
| Red | 269 | 0.13 | 0.42 |
| Yellow | 14,272 | 6.95 | 13.99 |
| Targets of concern (Red + Yellow) | 14,541 | 7.09 | 14.41 |
| Green | 1,686 | 0.82 | 2.13 |
| Grey (unclassified) | 188,978 | 92.09 | 83.47 |

**Table 3.** Total numbers of evaluated targets of concern and their final categories.

| Target Type | Total<br>Considered Based on<br>Preliminary Risk Tiers | Post-LLM Category Refinement |  |  |
| --- | --- | --- | --- | --- |
|  |  | Red | Yellow | Grey |
| Toxin targets | 2,039 | 70 | 1,060 | 909 |
| Viral receptors | 85 | 4 | 77 | 4 |
| Essential proteins | 9,130 | 171 | 6,898 | 2,061 |
| Immune system | 2,190 | 17 | 1,847 | 326 |
| Cell death pathways | 3,263 | 9 | 1,742 | 1,512 |
| Neurotransmitter systems | 998 | 20 | 646 | 332 |
| Coagulation system | 415 | 14 | 309 | 92 |
| Energy metabolism | 6,090 | 3 | 3,171 | 2,916 |

BLASTP achieved a higher F1 score (99.7% vs 97.2%) and area under the receiver operating characteristic curve (AUC, 99.7% vs 98.8%). This is not surprising for this evaluation, which tests a relatively simple case: detecting triple-mutation variants of database entries, where sequence identity provides a strong signal.

However, examining the 21 therapeutic targets that ESM-C incorrectly flagged revealed that most were members of the same protein family as a database entry, suggesting a confounding based on real, functional significance. For example, EPHA3 was flagged due to high cosine similarity (0.9956) with the dual-use target EPHA4; JAK1 with JAK2; CCR1 with CCR5; and three interferon-alpha subtypes (IFNA5, IFNA8, IFNA14) with IFNA2. All 21 matched yellow (dual-use) entries. BLASTP does not flag these evolutionarily related but sequence-divergent pairs, which accounts for its higher F1 score on this evaluation. Depending on the deployment context, it is possible that this additional flagging of evolutionarily and functionally similar proteins by the ESM-C method is desirable. This postulate on functional analysis is supported by the robustness analysis below.

### 2.3 Robustness to sequence variation

To evaluate robustness to sequence variation, we introduced increasing numbers of SNVs (10–100) into targets of concern and measured flagging rates. We analyzed benign and damaging mutations separately, as classified by PolyPhen predictions.^17^ For benign SNVs (Figure 4A), both the ESM-C embedding-space-based and BLASTP protein-sequence-based methods maintained high flagging rates at low mutation counts, but they diverged as mutations accumulated. At 100 benign SNVs, ESM-C flagged 51.4% of variants whilst BLASTP flagged only 9.6%. For damaging SNVs (Figure 4B), ESM-C showed markedly faster decay in flagging rate compared to benign SNVs; at 20 SNVs, ESM-C flagged only 54.6% of damaging variants compared to 97.0% of benign variants. In contrast, BLASTP showed nearly identical curves for benign and damaging SNVs (98.8% vs 98.8% at 20 SNVs; 43.6% vs 43.8% at 50 SNVs), consistent with the mechanism being insensitive to whether mutations disrupt protein function.

**Figure 4:**
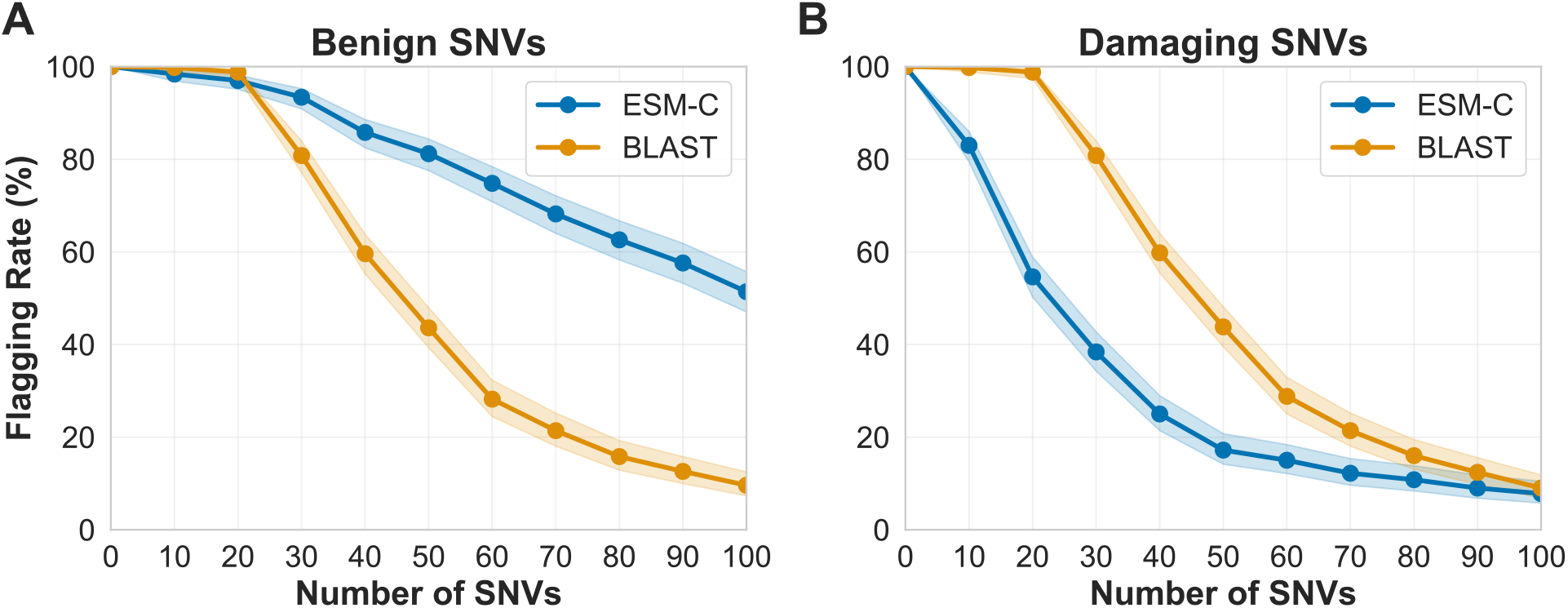
Robustness of screening methods to sequence variation. (A-B) Flagging rate as a function of the number of SNVs introduced into targets of concern for benign SNVs (A) and damaging SNVs (B). ESM-C (blue) and BLASTP (orange) were evaluated at their respective optimal thresholds. Shaded regions indicate 95% confidence intervals.

This difference reflects a fundamental distinction between the methods. ESM-C embeddings seem to capture structural and functional similarity, which would allow structurally disruptive mutations that reduce similarity to the wild-type to lead to markedly different flagging rates. By contrast, BLASTP measures sequence identity regardless of structural or functional impact. These findings suggest that ESM-C may capture structural and functional information, which supports the above postulate that a tool using the ESM-C may benefit biosecurity efforts where function of concern is the relevant metric over sequence similarity.

We also compared predicted structural similarity for a subset of variants; see Supplementary Figure S1.

### 2.4 Flagging rates across diverse protein datasets

To characterize how screening would affect different research contexts, we measured flagging rates across diverse datasets, including: proteins across a range of commonly studied species, proteins studied in general academic research, commercially relevant therapeutic targets, and experimentally validated binder targets.

To estimate expected flagging rates for researchers working with different organisms, we analyzed proteomes weighted by publication frequency (Figure 5A). We assessed human proteins as well as those from a range of species with varying evolutionary distance from humans, including mice, zebrafish, drosophila, fungi, plants, bacteria, and viruses. For each of these organisms we analyzed research-weighted samples of 2,000 reviewed Swiss-Prot proteins. They were drawn with sampling probability scaled by (log) PUBMED publication count, so the sample reflects the proteins researchers study, rather than the proteome at large.

**Figure 5:**
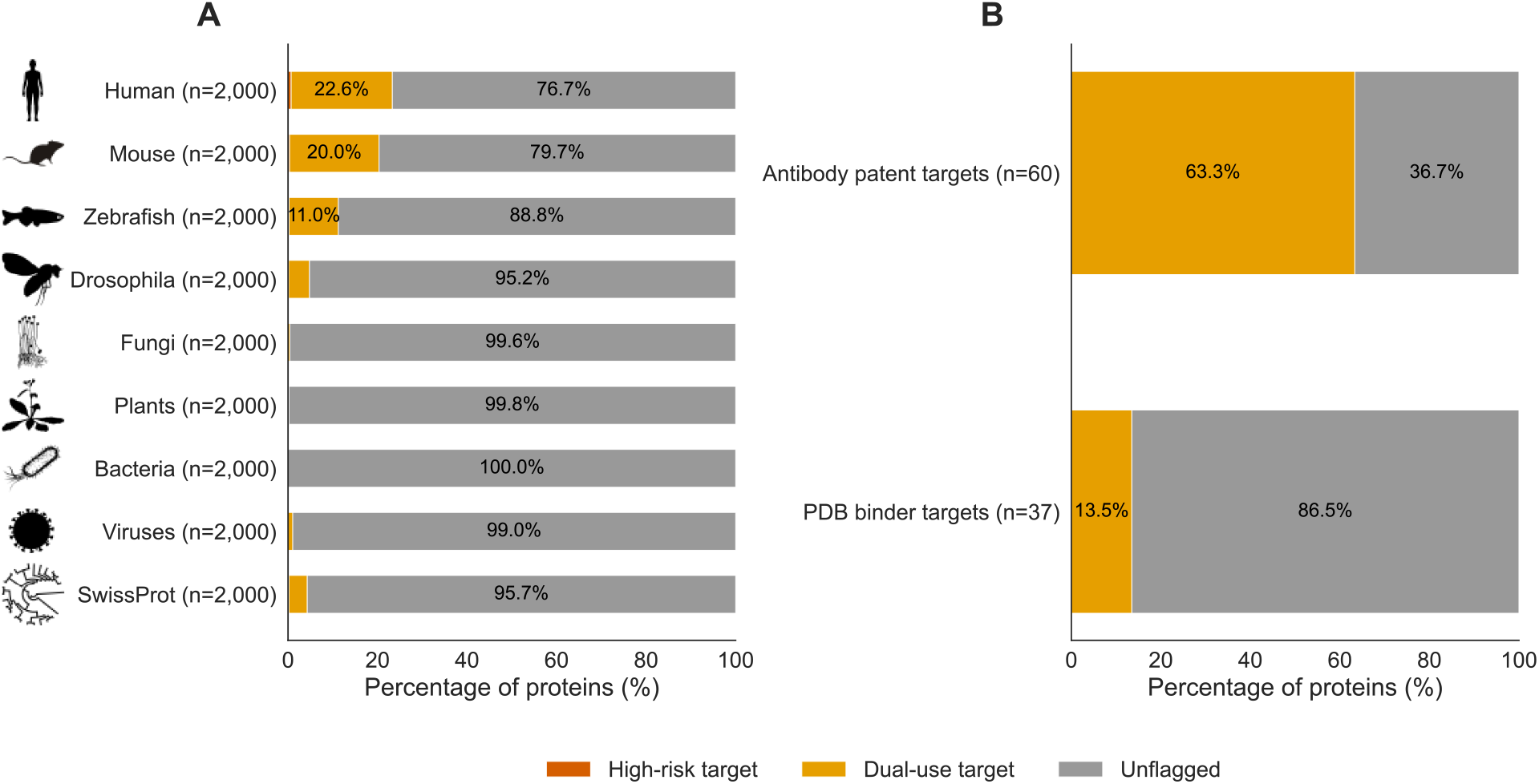
Flagging rates across diverse protein datasets. Proteins were screened against the targets of concern database using ESM-C embeddings at the optimized threshold (0.9903). Colors indicate risk classification: high-risk targets (red), dual-use targets (yellow), and unflagged (grey). (A) Cross-species proteomes sampled and weighted by research frequency (n=2,000 each). High-risk counts: human 14, mouse 7, zebrafish 5, Drosophila 3, fungi 1, all others 0. We also assessed a random sample from Swiss-Prot weighted by research frequency (n=2,000; 4 red, 82 yellow). (B) Protein design-relevant datasets: antibody patent targets from recent patent literature (n=60; 0 red, 38 yellow), and experimentally validated protein binder targets from the PDB (n=37; 0 red, 5 yellow).

For human proteins, 0.7% were flagged as high-risk (red category) and 22.6% as dual-use (yellow category), which amounted to 23.3% flagged total (14/2,000 high-risk, 452/2,000 dual-use). This exceeds the 7.1% of the human proteoforms classified as targets of concern, either high-risk or dual-use, during database construction (see methods section 4.1). The difference reflects three factors. The main factor is that research-weighted sampling over-represents well-studied proteins with critical biological functions, many of which are dual use. Additionally, reviewed Swiss-Prot proteins are better annotated than the full proteome and embedding-based screening captures functionally similar proteins whose annotations do not explicitly place them in concern categories.

The database contains only human targets of concern, so flagging of non-human proteins reflects embedding-space similarity to human targets of concern. Mouse proteins showed similar patterns with 0.35% flagged as high-risk and 20% as dual-use (20.4% flagged total: 7/2,000 high-risk, 400/2,000 dual-use), reflecting the high structural and functional conservation between mammalian orthologs. Flagging rates for model organisms with intermediate evolutionary distance to humans were lower: zebrafish (11.2% total flagged) and Drosophila (4.8% total flagged). For distantly related organisms, flagging rates dropped substantially on these research-weighted subsets: viruses (1.1% total flagged), fungi (0.4%), plants (0.2%), and bacteria (0%). These results suggest that the screening system should minimally disrupt research on organisms distantly related to humans, including work designing binders to microbial, plant, fungal, or viral proteins.

To estimate flagging rates for general academic research more broadly, we sampled all Swiss-Prot proteins weighted by publication frequency, providing a sample biased toward well-studied proteins. This research-weighted sample showed a flagging rate of 4.1% (82/2,000) as dual-use and 0.2% (4/2,000) as high-risk.

In addition to testing flagging rates for a range of species and for general academic research, we also analyzed several other types of protein targets that reflect plausible protein binder target use-cases within industry and academic research.

Antibody patent targets showed the highest flagging rate: 63.3% (38/60) were flagged as dual-use (yellow category), though none were flagged as high-risk (red category) (Figure 5B). These targets represent proteins of active commercial interest, and the high flagging rate reflects the substantial overlap between commercially valuable targets and biologically critical proteins. For example, CD4 was flagged as dual-use. CD4 is the receptor HIV uses to enter T cells; a designed binder could block viral entry (as the therapeutic antibody ibalizumab does), but could equally deplete or disable CD4^+^ T cells, causing immunodeficiency.^18^ This illustrates the inherent dual-use nature of many therapeutically important proteins and highlights the potential value of using a tiered managed access system in concert with this tool, as addressed in the discussion.

For experimentally validated protein binder targets deposited in the PDB, fewer proteins were flagged: 13.5% (5/37) as dual-use, and none as high-risk. Although this database is small, these proteins best reflect the real-world implications of our screening methods as they are proteins for which scientists have created binders.

## 3 Discussion

### Contributions

We propose the first screening framework designed for implementation across an entire class of biological AI models, using protein binder design tools as an initial application. We catalogued 14,541 human proteins with essential functions that would be concerning if disrupted, assigning risk levels based on dual-use potential. We developed a screening method using protein language model embeddings that captures functional relationships between proteins, enabling detection of targets of concern even when sequences differ due to natural variation.

Our evaluation demonstrated that ESM-C embedding-based screening reliably identifies variants of targets of concern while allowing unrelated therapeutic targets to pass. Critically, ESM-C discriminates between benign mutations–which preserve protein function and should be flagged–and damaging mutations–which likely disrupt function and need not be flagged. This sensitivity to functional impact distinguishes embedding-based screening from sequence alignment methods, which treat all mutations equivalently regardless of their biological consequences.

We used LLM-based assessment to refine risk classifications, enabling systematic evaluation of whether targeting each protein would actually cause harm. This reduced the database from 22,299 to 14,541 targets of concern by filtering out proteins where functional annotations suggested concern, but biological analysis indicated low risk. We anticipate that this database will need to be periodically updated to incorporate new information about hazards and therapeutic utility.

We characterized how screening would affect different research contexts. For organisms closely related to humans, flagging rates are significant (23% for human, 20% for mouse), reflecting the high conservation of essential proteins across mammals. Flagging rates decline with evolutionary distance: 11% for zebrafish, 5% for Drosophila, and below 1.1% for bacteria, fungi, plants, and viruses. This pattern is expected, since the database contains only human targets of concern and sequence similarity to human proteins decreases with phylogenetic distance. Researchers working on distantly related organisms should experience minimal disruption. Among commercially relevant targets, 63% of antibody patent targets were flagged as dual-use, reflecting that therapeutically important proteins often perform critical biological functions.

### Limitations

The database is currently limited to human proteins. Expansion to other species, such as key crop species or livestock, would increase coverage but also increase flagging rates for legitimate agricultural research.

We optimized thresholds on an evaluation dataset of SNV and therapeutic targets. Real-world performance may differ, underscoring the importance of monitoring-first deployments to gather calibration data. The evaluation also focused on SNVs with high sequence identity to database entries; performance on more divergent sequences, such as distant homologs or sequences deliberately modified to evade detection, remains to be characterized.

### Policy Implications

The significant overlap between therapeutic targets and targets of concern means that many legitimate research queries are likely to be flagged. This reflects genuine biological reality: proteins with critical functions are often both therapeutically important and potentially dangerous if disrupted.

Additional refinement of the screening reference database can likely streamline the high-risk and dual-use categories in this screening approach to reduce flagging rates without missing genuinely risky user requests. However, significant overlap is very likely to persist.

To address this challenge, deployment of user-input screening approaches will need to be done in concert with other complementary biosecurity interventions. Options to consider include holding flagged requests for follow-up review as opposed to immediate blocking, and/or tiered responses rather than uniform blocking. It will also be important to explore integration of user input screening with managed-access deployment of protein design models, where it would be possible to fulfill dual-use requests to trusted users that have undergone more significant know-your-customer screening.

### Future work

One of the most important next steps is real-world deployment and evaluation of this screening approach. Releasing the screening software system as an open-source Python package communicating with a secure API that holds the reference database would enable tool developers to integrate screening with minimal friction. Initial deployments could use monitoring mode to gather data on typical usage patterns and flag rates, informing refinements to the database and threshold before any blocking is considered.

For tools available only to vetted users, such as Chai-2^7^, a protein binder design tool accessible only through Chai Discovery’s Responsible Deployment policy, screening can complement existing access controls and know-your-customer practices. Flagged requests could trigger enhanced logging or periodic review rather than refusal, allowing legitimate research to proceed while maintaining oversight. Screening could function effectively as one layer in a defense-in-depth approach that includes access controls, user vetting, and community norms.

Further evaluation of this screening approach should test robustness to deliberate evasion attempts. For example, we tested whether appending a disordered tail to a target of concern evades screening (Supplementary Figure S2). It does for 99 of 100 targets, because averaging the embedding over the whole protein dilutes the match, but screening the request in short overlapping windows recovers detection. Developing windowed screening into a robust defense, and red-teaming the screen against other evasions, is important future work.

Alternative screening approaches could also be explored; for example, structure prediction methods such as ESMFold may better capture functional properties but would introduce additional computational overhead. Protein language model embeddings offer a practical middle ground, providing stronger functional sensitivity than sequence alignment while remaining computationally efficient.

Screening can and should extend to other biological AI tool categories. A natural next step would be protein design tools more broadly, which may require output screening rather than input screening. The embedding-based approach may also be relevant for DNA synthesis screening, where it could complement existing sequence alignment methods without requiring model fine-tuning.

### Conclusion

We present a novel approach for screening user inputs to protein binder design tools. The goal of this work has been to provide an early proof of principle that can be refined and generalized to other protein design tools and broader categories of biological AI models. It is important to explore new solutions in this area because BAIMs are likely to introduce new biosecurity risks, and there has been limited development and deployment of guardrails for BAIMs.

Our assessment shows that this protein-langauge-model-based approach performs better than BLASTP at identifying binder targets that have functional similarity but lower sequence homology, and it offers a path to transcending traditional sequence-based approaches to screening. In an era where BAIMs are designing constructs that have low homology to known sequences, this will be vital for continued advances in biosecurity.

Importantly, this screening approach is designed to identify and flag risky user requests from protein binder design tools without placing an undue burden on scientific research and innovation. To achieve this balance, it will be essential to deploy this screening approach in a way that addresses the overlap our analysis showed between targets of concern and therapeutic targets. Effective deployment will require further refinement of the screening database to reduce false positives, while also integrating screening with broader governance structures that allow for tiered responses, managed access approaches, know-your-customer vetting, usage monitoring, and clear policies for responding to flags. For the research community more broadly, it means developing shared norms around responsible use of biological AI tools and contributing to ongoing refinement of screening databases and methods. Screening alone cannot prevent misuse, but as part of a layered approach, it can meaningfully reduce risk while preserving the substantial benefits of biological AI for legitimate research.

## 4 Methods

### 4.1 Database of targets of concern

#### 4.1.1 Sources

We downloaded the human reference proteome from UniProt (*Homo sapiens*, NCBI taxonomy ID 9606) on August 27th, 2025. It contains 205,205 protein entries: 20,420 reviewed (Swiss-Prot) and 184,785 unreviewed (TrEMBL). We used the full set, including unreviewed entries, to classify human targets of concern as comprehensively as possible.

We defined eight categories. For each, we assembled a reference set of human proteins from a curated source and labelled every proteome entry that matched that reference set. Matching was by shared UniProt accession, by gene symbol, or by Gene Ontology annotation, depending on the source. *Note: the specific sources are omitted for biosecurity reasons*.

- **Toxin targets (red):** human proteins that known toxins act on; a binder could reproduce a toxin’s effect. Matched by shared UniProt accession ID
- **Viral receptors (red):** host proteins that viruses use to enter cells; a binder could act as a novel viral entry protein. Matched by shared UniProt accession ID
- **Essential proteins (yellow):** proteins required for cell survival. Matched by gene symbol
- **Immune system proteins (yellow):** proteins whose disruption causes immunodeficiency. Matched by gene symbol
- **Cell death pathways (yellow):** identified by Gene Ontology annotation
- **Neurotransmitter systems (yellow):** targets of nerve agents and neurotoxins. Identified by Gene Ontology annotation
- **Coagulation system (yellow):** proteins whose disruption causes bleeding or thrombosis. Identified by Gene Ontology annotation
- **Energy metabolism (yellow):** core energy-production pathways. Identified by Gene Ontology annotation

For the Gene Ontology categories, we counted a protein as a match if its UniProt GO annotations contained the target term or any of its descendant terms retrieved from the Gene Ontology. Annotations propagate across isoforms: every entry sharing a gene with a flagged protein receives the same label, which is why the counts are at the proteoform level rather than the gene level.

#### 4.1.2 Preliminary risk tiers

Each candidate was assigned a preliminary risk tier. Toxin targets and viral receptors were categorized as high risk (red): they have the clearest offensive purpose, since a binder to them could directly enable harm, and they have few legitimate uses. The other six categories were categorized as dual-use (yellow): disrupting them can be harmful, but they also have clear research and therapeutic value.

Separately, we labeled clinical drug targets as a benign set (green), used only as the negative set for the analyses, not as part of the screening database. We removed from the green set any protein that is an isoform of, or near-identical in sequence to, a target of concern, so the benign and concern sets do not overlap.

When a target was flagged in multiple different categories, it was classified as the highest-risk tier. If it was flagged in both the highest risk tier (red) and as a clinical drug target, it was classified as dual use (yellow) because it has both harmful and beneficial uses.

These tiers were provisional. As described in the next section, we refined the preliminary risk tiers using a large language model.

### Refinement of preliminary risk tiers using a language model

The categories described above tend to over-flag: protein targets are defined as candidates for the high-risk (red) category or dual-use (yellow) category, regardless of whether an engineered binder to them would actually be harmful or tractable. We used a language model to refine the classification, scoring each candidate and assigning its final tier.

We graded every protein target, which was initially designated in the red or yellow categories, with Claude Sonnet 4.6 (model ID claude-sonnet-4-6). For each protein we supplied its UniProt annotations (function, subcellular location, tissue expression, disease involvement, and pharmaceutical use) and requested two scores from 1 to 5 in two categories:

- **Harm:** Scoring from 1-5 based on the worst-case consequence of an engineered binder engaging the target. The model considered all plausible binding outcomes (blocking, activating, or crosslinking the protein) and scored the most harmful, weighing mechanism, severity, how many tissues are affected, onset, and redundancy.
- **Benefit:** Scoring from 1-5 of the protein’s legitimate therapeutic value.

Because intracellular targets are harder to reach with a binder, we applied a fixed accessibility discount to the harm score by subcellular location: extracellular and cell-surface proteins took no discount, cytoplasmic and nuclear proteins minus one, and mitochondrial proteins minus two. The model reported both the raw and discounted scores.

We mapped the two scores to a final tier deterministically. Binding targets with a harm score of 2 or lower were classified as grey and removed from the database. Targets were assigned to the yellow dual-use category if they had a high harm score of 4 or 5 coupled with a benefit score of 3-5, or if they had a harm score of 3. Targets were assigned to the red category if they had a high harm score of 4 or 5 with benefit of 2 or below. Grades propagate across isoforms sharing a gene.

The model graded 97.1% of the targets of concern (14,120 of 14,541). The remaining 2.9% (421) had empty UniProt function fields or were unreviewed and therefore could not be graded; these retained their preliminary risk tier described above. After refinement the database contained 14,541 targets of concern (269 red, 14,272 yellow), down from 22,299.

#### 4.1.3 Validation of the grading

We audited the grading through manual review of approximately 50 gradings, sampled deliberately to stress-test the model: a spread of moderate harm (score 3) proteins across all eight categories, and a sample of the highest-harm (harm 4 and 5) proteins, and every case where the model’s own recommendation disagreed with the deterministic scoring rule. For each we checked that the harm and benefit reasoning was sound.

The audit surfaced one systematic error: the model under-scored the harm of viral receptor targets, because it did not account for their potential to create transmissible agents (a binder acting as a novel viral entry protein). We corrected the three affected viral receptors (CLEC4G, MFSD6, ITGB6). We also corrected one essential protein (GARS1), where the intracellular-accessibility discount had been over-applied to an enzyme with a secreted fraction. These four manual overrides were applied last in the pipeline.

#### 4.1.4 Summary table: post-LLM refinement of risk tiers

We report each count as a fraction of the full proteome (205,205 entries) and of the reviewed proteome (20,420 reviewed entries): % of proteome = count / 205,205; % of reviewed = reviewed proteins in the row / 20,420.

Targets of concern are 7.1% of the whole human proteome but 14.4% of the reviewed proteome. Reviewed (better-studied) proteins are enriched for targets of concern, which is why the flagged fraction is roughly twice as high among them.

Each protein target type was graded as red, yellow, or grey categories. Proteins can belong to more than one category, so category totals overlap; within a row, red + yellow + grey sum to the total.

### 4.2 Screening methods

We developed an embedding-based screening method using ESM-C, a protein language model that generates superior protein representations compared to larger ESM-2 models while maintaining computational efficiency.^19^ To benchmark this approach, we compared it to BLASTP, the established sequence-based alignment-based method.^12^ We ran BLAST+ v2.17.0 throughout with default scoring (BLOSUM62, gap open 11, extend 1, E-value 10, word size 3).

For ESM-C screening, we generated embeddings by encoding each sequence through the model and mean-pooling hidden states across sequence length to produce a fixed 1,152-dimensional representation. We pre-computed embeddings for all 14,541 targets of concern in the database. For each query sequence, we computed cosine similarity to all targets of concern embeddings and recorded the maximum similarity score. We flagged sequences where the maximum similarity exceeded the optimized threshold of 0.9903.

For BLASTP screening, we constructed a BLASTP database from the 14,541 target of concern protein sequences.^13^ We ran BLASTP for each query sequence against this database and recorded the single best hit. We used percent sequence identity of the top hit as the similarity score, normalized to a 0–1 scale. We flagged sequences where sequence identity exceeded the optimized threshold of 0.9397 (93.97%).

Both methods were evaluated using identical datasets and threshold optimization procedures to ensure fair comparison.

### 4.3 Variant retrieval and classification

The following threshold optimization and robustness analyses used annotated natural variants of targets of concern. We downloaded all missense variants from the EBI Proteins API, retrieving 2,631,615 mutations across 4,424 targets of concern with available data.^20^ We classified mutations using PolyPhen predictions from the EBI API.^17^ We categorized mutations as benign if PolyPhen predicted “benign” and as damaging if PolyPhen predicted “probably damaging.” We excluded mutations classified as “possibly damaging” (low confidence) and those without PolyPhen predictions, comprising 33.3% of total mutations. Final classification yielded 1,110,547 benign mutations (42.2%) and 644,857 damaging mutations (24.5%).

### 4.4 Threshold optimization and method comparison

We constructed an evaluation dataset with two classes: sequences that should be flagged (variants of targets of concern) and sequences that should pass (therapeutic targets with no direct overlap to targets of concern).

To generate should-flag examples, we applied benign mutations to targets of concern. We generated triple-SNVs by randomly sampling and applying three mutations per target of concern, yielding 3,625 variants from 3,625 unique targets of concern. We chose three SNVs to encompass the average variation of two SNVs found in human proteins,^21^ plus a conservative buffer.

For should-pass examples, we used clinically validated therapeutic targets. We obtained therapeutic targets from the Open Targets Known Drug dataset, which contains target gene products for approved or clinical candidate drugs derived from ChEMBL.^22^ To ensure clean separation from targets of concern, we removed therapeutic targets if they shared gene names with any target of concern (including isoforms) or had greater than 80% Jaccard sequence similarity to any target of concern.

We split the dataset into calibration (70%) and test (30%) sets. The split was performed at the target of concern level for should-flag examples to prevent data leakage, ensuring all variants from the same target of concern appear in the same split. Final dataset sizes were: calibration set with 2,520 target of concern variants and 1,174 therapeutic targets; test set with 1,081 target of concern variants and 504 therapeutic targets.

For threshold optimization, we tested 200 candidate thresholds across the observed similarity range for each method. For each threshold, we classified sequences as flagged (similarity at or above threshold) or passed (similarity below threshold) and computed precision, recall, and F1 score on the calibration set. We selected the threshold maximizing F1 score on the calibration set and evaluated final performance on the held-out test set. We computed the area under the receiver operating characteristic curve (AUC) on the test set.

To confirm the F1 and AUC gap was not an artifact of one split, we repeated calibration and evaluation over 100 random 70/30 splits, grouped by source target of concern so no target appeared in both folds. Each split returned the threshold to maximize F1 on the calibration fold, scored the held-out fold, and used the same split for both methods (paired). We tested the mean paired difference with the Nadeau and Bengio corrected resampled t-test,^23^ which accounts for the overlap between calibration folds and checked the direction with a Wilcoxon signed-rank test. Both methods stayed strong (ESM-C F1 96.9 ± 0.3%, BLASTP 99.6 ± 0.1%), the ESM-C threshold was stable (0.9903 ± 0.0004), and the small gap favoring BLASTP was significant for both F1 and AUC (p < 0.001).

We also tested sensitivity to class imbalance. The evaluation set has about twice as many targets of concern as therapeutic targets (68:32). Subsampling the targets of concern to a 50:50 balance and re-optimizing raised the ESM-C threshold only slightly, from 0.9903 to 0.9939 (more stringent, so marginally fewer flags). We retained 0.9903, optimized on the full evaluation set, throughout.

### 4.5 Robustness to sequence variation

To evaluate how screening performance degrades with increasing sequence divergence, we generated mutant sequences with varying numbers of mutations and measured flagging rates.

We randomly selected 500 targets of concern with at least 100 benign and 100 damaging mutations to ensure balanced sampling across all mutation counts. For each target of concern, we generated mutant sequences with 10, 20, 30, 40, 50, 60, 70, 80, 90, and 100 mutations for both benign and damaging categories. We sampled mutations randomly without replacement from the available pool for each category. This produced 10,000 mutant sequences in total (500 targets of concern × 10 mutation counts × 2 mutation types). We screened all mutant sequences using both ESM-C and BLASTP at their respective optimized thresholds.

To investigate the relationship between sequence identity, structural similarity, and flagging decisions, we predicted structures for a subset of mutants. We focused on benign mutations only, as these represent sequences that should retain function and be flagged. We filtered to 30 targets of concern with sequences of 400 amino acids or fewer due to ESMFold API length constraints. For mutation counts of 10, 20, 50, and 100, we computed sequence identity between each mutant and its wild-type as the fraction of identical residues. We predicted structures for both wild-type targets of concern and mutant sequences using the ESMFold API.^2^ We computed structural similarity using TM-score via TM-align, normalized by wild-type sequence length.^24^ The final structural analysis included 240 mutant-wildtype pairs (30 targets of concern × 4 mutation counts × 2 replicates).

### 4.6 Flagging rates across diverse protein datasets

To characterize how screening would affect different research contexts, we screened proteins from diverse datasets.

For antibody patent targets, we extracted unique target names from a recent survey of antibody patents (Peng et al., 2025, Supplementary Data 1).^25^ For each unique target name, we queried UniProt for reviewed human proteins matching the gene name.^26^ We mapped 60 of 69 unique targets to UniProt sequences; the remaining 9 had no matching canonical protein entry.

For PDB binder targets, we queried RCSB PDB using full-text search for “designed binder” across all experimental structures.^27^ For each structure, we retrieved FASTA sequences and extracted natural target proteins by filtering out synthetic constructs (NCBI taxonomy ID 32630) and proteins with names containing design-related keywords (“binder”, “designed”, “de novo”, “engineered”, “artificial”, “synthetic”). This yielded 37 unique target sequences from structures containing designed protein binders.

To generate a research-weighted Swiss-Prot sample representative of proteins that researchers actually work with, we weighted sampling by publication frequency. We sampled an initial pool of 20,000 random Swiss-Prot accessions and retrieved gene names for each protein from UniProt.^26^ We queried PubMed for each gene name using the NCBI E-utilities API. We excluded gene names shorter than 3 characters, which return inflated counts due to matching common abbreviations. We performed weighted random sampling without replacement, using log-transformed publication counts as weights to balance representation of well-studied proteins without extreme skew. The final sample size was 2,000 proteins.

For cross-species analysis, we applied the same research-weighted sampling approach to proteomes from six organism groups, using NCBI taxonomy IDs to filter reviewed Swiss-Prot entries: human (9606), mouse (10090), zebrafish (7955), Drosophila (7227), bacteria (2), plants (33090), fungi (4751), and viruses (10239).^26^ For each group, we sampled an initial pool of up to 20,000 proteins, queried PubMed counts, and performed weighted sampling to obtain 2,000 proteins per group.

We screened all proteins using ESM-C at the optimized threshold of 0.9903.^16^ The flag level (high-risk or dual-use) was determined by the risk classification of the matched target of concern.

## 5 Data and code availability

Code for this project is available at: https://github.com/PhilPalmer/bioscreen. The screening database and supplementary information on database construction are available upon request for researchers with a legitimate need, subject to biosecurity review. Parties interested in the screening database are encouraged to contact NTI | bio.

## 6 Acknowledgements

The authors thank James Diggans, Anthony Gitter, Richard Moulange, Ryan Ritterson, and Sana Zakaria for their thoughtful review and feedback on this manuscript and earlier technical specifications, which substantially improved this work.

This work was commissioned by the Nuclear Threat Initiative Global Biological Policy and Programs (NTI | bio) in support of the AIxBio Global Forum, with generous funding from Sentinel Bio and Coefficient Giving.

## 7. Supplementary Figures

The scatter plots (Figure S1A-B) reveal the decision boundaries of each method. ESM-C (Figure S1A) primarily flags sequences with high sequence identity (>90%), though some lower-identity sequences are also flagged. Surprisingly, the TM-score (indicating structural similarity) does not appear to be strongly correlated with flagging decisions. BLASTP (Figure S1B) shows a sharp threshold around 94% sequence identity, consistent with the optimized threshold of 93.97%.

In our structural analysis of 240 protein pairs, we did not observe a clear relationship between structural similarity and flagging decisions. Whether this reflects a genuine property of the embedding space or limitations of our analysis remains an open question. TM-scores reflect global backbone geometric similarity and may score highly due to the widespread reuse of common folds like the TIM barrel and Rossmann domain across functionally unrelated enzyme classes (Holm & Sander, 1996; Nagano et al., 2002). TM-scores are largely insensitive to the local sequence and structural features such as active site composition, surface electrostatics, and loop conformations that actually determine protein function. Given these limitations, it is not clear whether TM-scores or ESM-C-embedding-space similarity are more representative of ground truth.

**Figure S1:**
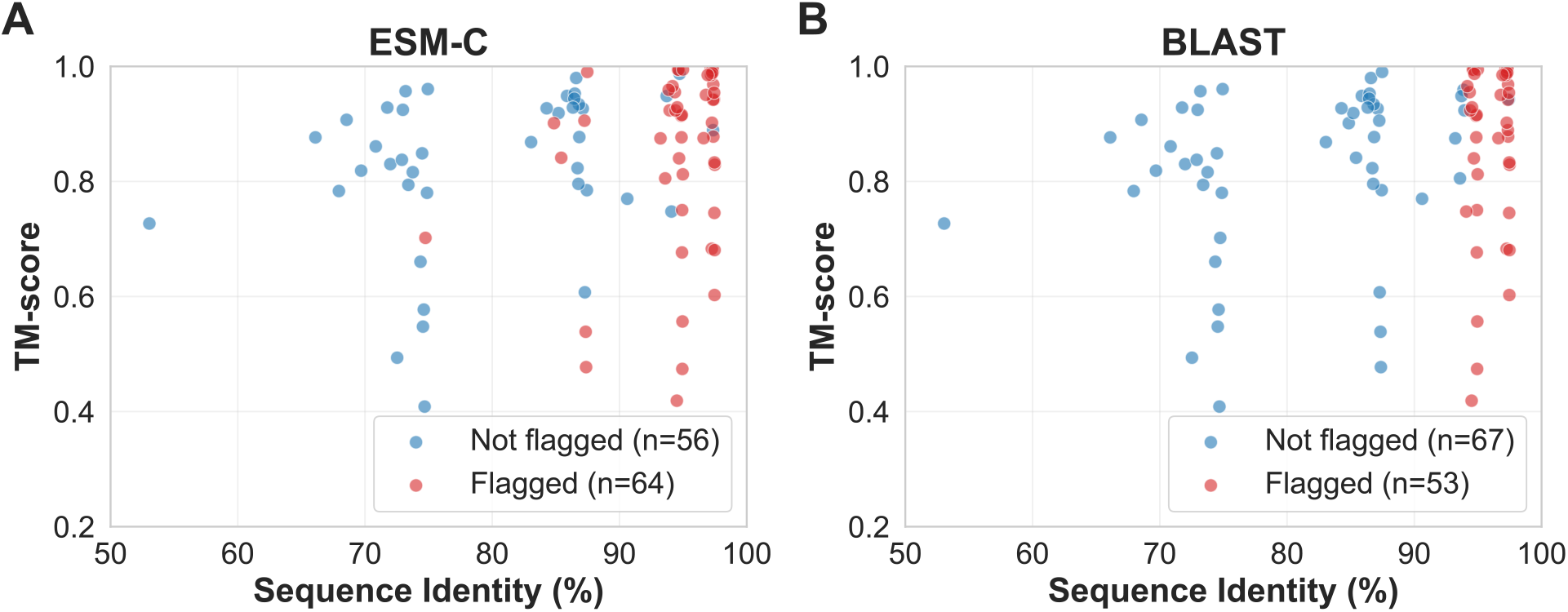
Sequence identity versus predicted structural similarity for flagged and unflagged variants. Sequence identity to the nearest database target (x-axis) against predicted structural similarity to wild-type (TM-score, y-axis) for ESM-C (A) and BLASTP (B) screening. Points are colored by flagging status (red = flagged, blue = not flagged). The analysis is restricted to targets of concern with predicted-benign SNVPs (PolyPhen^17^) and sequences 400 amino acids due to ESMFold^2^ API length limits (n=240 pairs); this subset is length-biased and not a random sample. TM-scores were computed with TM-align^24^ and reflect global backbone geometry only, so they may miss the local features that determine function and are not a definitive ground truth for functional similarity.

Robustness to disordered-tail extension. We tested whether appending a disordered region to a target of concern can evade embedding screening while leaving the folded domain intact. For each of 100 distinct targets of concern (one representative entry per gene, 450 to 900 residues; 32 high-risk and 68 dual-use), we built a construct consisting of the target sequence, a 20-residue glycine-serine linker (GGGGSGGGGSGGGGSGGGGS), and a 300-residue disordered region from BRCA1 (UniProt P38398). We screened each construct as a whole protein (mean-pooled ESM-C embedding) against the targets-of-concern database, and separately in overlapping 400-residue windows under two schemes: comparing each request window to the whole-protein database and comparing each request window to a windowed version of the database. A construct evaded if its whole-protein similarity fell below the 0.9903 threshold and was recovered if the best window reached the threshold.

Ninety-nine of the 100 constructs evaded whole-protein screening: appending roughly 100 or more disordered residues dilutes the mean-pooled embedding below the threshold, while the 20-residue linker alone does not (Figure S2A). Screening in windows recovered the target, but recovery depended on how the database was represented (Figure S2B). Comparing request windows to the current whole-protein database recovered fewer than half of the targets (46/100). Comparing request windows to a windowed database recovered all of them (100/100, similarity 1.0), because a window over the unchanged folded domain is an exact copy of a database window. This holds only while the domain is unmodified; a mutated domain would not match, which is the separate robustness axis tested in the main text. We did not measure the false-positive cost of windowed screening and used a single 400-residue window on proteins of 450 to 900 residues, so windowed screening is a direction for future work rather than a validated defense.

**Figure S2:**
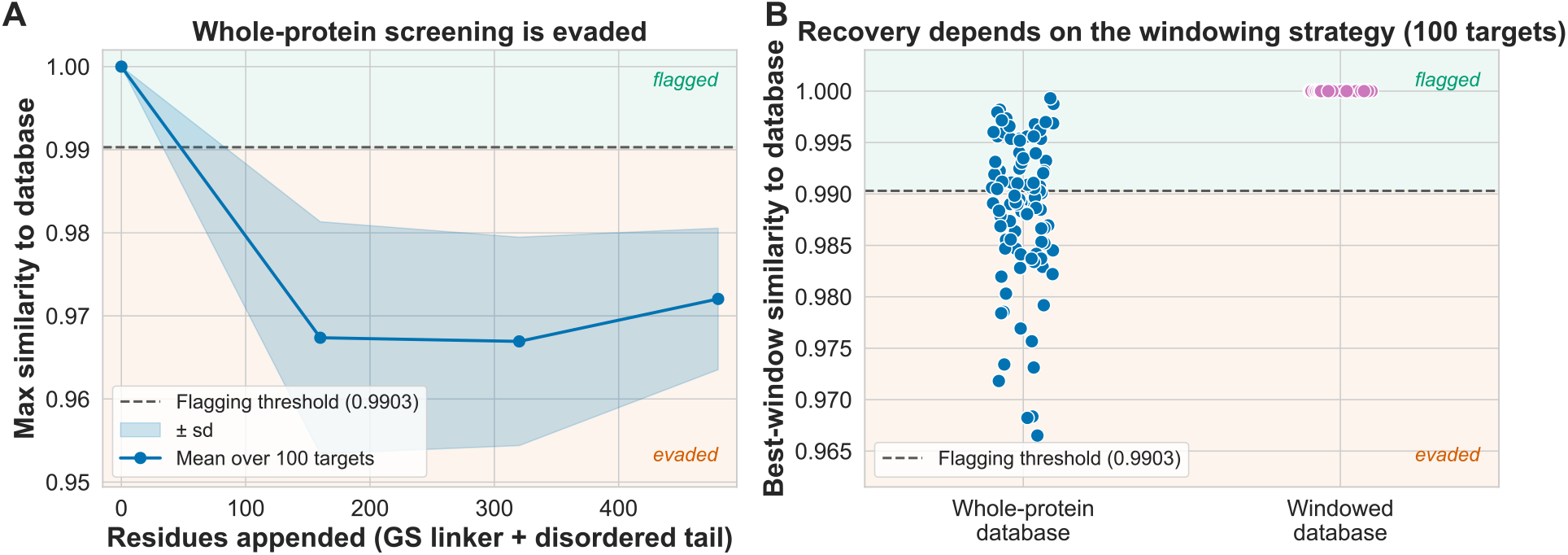
Evasion by disordered-tail extension, and recovery by windowed screening. (A) Appending a glycine-serine linker and an increasing disordered tail to a target of concern lowers its maximum cosine similarity to the targets-of-concern database below the flagging threshold (0.9903). The line and shaded band show the mean ± standard deviation across 100 distinct targets of concern; 99/100 evade. (B) Recovery of the same constructs when screened in 400-residue windows, one point per target: comparing each window to the whole-protein database recovers 46/100, whereas comparing to a windowed version of the database recovers 100/100 at a similarity of 1.0. Points above the threshold are flagged; points below evade.

## References

[1] Josh Abramson et al. “Accurate structure prediction of biomolecular interactions with AlphaFold 3”. In: Nature 630.8016 (June 2024), pp. 493–500. ISSN: 1476-4687. DOI: 10.1038/s41586-024-07487-w. URL: https://www.nature.com/articles/s41586-024-07487-w (visited on 12/21/2025).

[2] Zeming Lin et al. “Evolutionary-scale prediction of atomic-level protein structure with a language model”. In: Science 379.6637 (Mar. 17, 2023), pp. 1123–1130. DOI: 10.1126/science.ade2574. URL: https://www.science.org/doi/10.1126/science.ade2574 (visited on 12/21/2025).

[3] Thomas Hayes et al. “Simulating 500 million years of evolution with a language model”. In: Science 387.6736 (Feb. 21, 2025), pp. 850–858. DOI: 10.1126/science.ads0018. URL: https://www.science.org/doi/10.1126/science.ads0018 (visited on 12/21/2025).

[4] Vinicius Zambaldi et al. De novo design of high-affinity protein binders with AlphaProteo. Sept. 12, 2024. DOI: 10.48550/arXiv.2409.08022. arXiv: 2409.08022[q-bio]. URL: http://arxiv.org/abs/2409.08022 (visited on 12/21/2025).

[5] Martin Pacesa et al. “One-shot design of functional protein binders with BindCraft”. In: Nature 646.8084 (Oct. 2025), pp. 483–492. ISSN: 1476-4687. DOI: 10.1038/s41586-025-09429-6. URL: https://www.nature.com/articles/s41586-025-09429-6 (visited on 12/10/2025).

[6] Hannes Stark et al. BoltzGen: Toward Universal Binder Design. ISSN: 2692-8205 Pages: 2025.11.20.689494 Section: New Results. Nov. 24, 2025. DOI: 10.1101/2025.11.20.689494. URL: https://www.biorxiv.org/content/10.1101/2025.11.20.689494v1 (visited on 12/21/2025).

[7] Chai Discovery Team et al. Zero-shot antibody design in a 24-well plate. ISSN: 2692-8205 Pages: 2025.07.05.663018 Section: New Results. July 6, 2025. DOI: 10.1101/2025.07.05.663018. URL: https://www.biorxiv.org/content/10.1101/2025.07.05.663018v1 (visited on 12/22/2025).

[8] Qiuzhen Li, Efstathios Nikolaos Vlachos, and Patrick Bryant. “Design of linear and cyclic peptide binders from protein sequence information”. In: Communications Chemistry 8.1 (July 22, 2025), p. 211. ISSN: 2399-3669. DOI: 10.1038/s42004-025-01601-3. URL: https://www.nature.com/articles/s42004-025-01601-3 (visited on 12/21/2025).

[9] Nathaniel R. Bennett et al. “Atomically accurate de novo design of antibodies with RFdiffusion”. In: Nature (Nov. 5, 2025), pp. 1–11. ISSN: 1476-4687. DOI: 10.1038/s41586-025-09721-5. URL: https://www.nature.com/articles/s41586-025-09721-5 (visited on 12/21/2025).

[10] Joseph L. Watson et al. “De novo design of protein structure and function with RFdiffusion”. In: Nature 620.7976 (Aug. 2023), pp. 1089–1100. ISSN: 1476-4687. DOI: 10.1038/s41586-023-06415-8. URL: https://www.nature.com/articles/s41586-023-06415-8 (visited on 12/21/2025).

[11] David Atanasov, Niccolò Zanichelli and Jean-Stanislas Denain. Expanding our analysis of biological AI models. Epoch AI. URL: https://epoch.ai/blog/expanding-our-analysis-of-biological-ai-models (visited on 03/30/2026).

[12] “IGSC Harmonized Screening Protocol v3.0”. In: ().

[13] Christiam Camacho et al. “BLAST+: architecture and applications”. In: BMC bioinformatics 10 (Dec. 15, 2009), p. 421. ISSN: 1471-2105. DOI: 10.1186/1471-2105-10-421.

[14] Bruce J. Wittmann et al. “Strengthening nucleic acid biosecurity screening against generative protein design tools”. In: Science (New York, N.Y.) 390.6768 (Oct. 2, 2025), pp. 82–87. ISSN: 1095-9203. DOI: 10.1126/science.adu8578.

[15] Gary Abel et al. Beyond Sequence Similarity: Toward Function-Based Screening of Nucleic Acid Synthesis. Rochester, NY, Mar. 17, 2026. DOI: 10.2139/ssrn.6444478. URL: https://papers.ssrn.com/abstract=6444478 (visited on 03/30/2026).

[16] Evolutionary Scale · ESM Cambrian: Revealing the mysteries of proteins with unsupervised learning. URL: https://www.evolutionaryscale.ai/blog/esm-cambrian (visited on 12/21/2025).

[17] Ivan A. Adzhubei et al. “A method and server for predicting damaging missense mutations”. In: Nature Methods 7.4 (Apr. 2010), pp. 248–249. ISSN: 1548-7105. DOI: 10.1038/nmeth0410-248. URL: https://www.nature.com/articles/nmeth0410-248 (visited on 12/21/2025).

[18] Simona A. Iacob and Diana G. Iacob. “Ibalizumab Targeting CD4 Receptors, An Emerging Molecule in HIV Therapy”. In: Frontiers in Microbiology 8 (2017), p. 2323. ISSN: 1664-302X. DOI: 10.3389/fmicb.2017.02323.

[19] Luiz C. Vieira, Morgan L. Handojo, and Claus O. Wilke. “Medium-sized protein language models perform well at transfer learning on realistic datasets”. In: Scientific Reports 15.1 (July 1, 2025), p. 21400. ISSN: 2045-2322. DOI: 10.1038/s41598-025-05674-x. URL: https://www.nature.com/articles/s41598-025-05674-x (visited on 12/21/2025).

[20] Andrew Nightingale et al. “The Proteins API: accessing key integrated protein and genome information”. In: Nucleic Acids Research 45 (W1 July 3, 2017), W539–W544. ISSN: 0305-1048. DOI: 10.1093/nar/gkx237. URL: https://doi.org/10.1093/nar/gkx237 (visited on 12/21/2025).

[21] Peng Yue and John Moult. “Identification and Analysis of Deleterious Human SNPs”. In: Journal of Molecular Biology 356.5 (Mar. 10, 2006), pp. 1263–1274. ISSN: 0022-2836. DOI: 10.1016/j.jmb.2005.12.025. URL: https://www.sciencedirect.com/science/article/pii/S0022283605015871 (visited on 12/17/2025).

[22] Annalisa Buniello et al. “Open Targets Platform: facilitating therapeutic hypotheses building in drug discovery”. In: Nucleic Acids Research 53 (D1 Jan. 6, 2025), pp. D1467–D1475. ISSN: 1362-4962. DOI: 10.1093/nar/gkae1128. URL: https://doi.org/10.1093/nar/gkae1128 (visited on 12/17/2025).

[23] Claude Nadeau and Yoshua Bengio. “Inference for the Generalization Error”. In: Machine Learning 52.3 (Sept. 1, 2003), pp. 239–281. ISSN: 1573-0565. DOI: 10.1023/A:1024068626366. URL: https://doi.org/10.1023/A:1024068626366 (visited on 07/06/2026).

[24] Yang Zhang and Jeffrey Skolnick. “TM-align: a protein structure alignment algorithm based on the TM-score”. In: Nucleic Acids Research 33.7 (Apr. 1, 2005), pp. 2302–2309. ISSN: 0305-1048. DOI: 10.1093/nar/gki524. URL: https://doi.org/10.1093/nar/gki524 (visited on 12/21/2025).

[25] Y. Peng et al. “Monoclonal antibody formulations: a quantitative analysis of marketed products and patents”. In: mAbs 17 (2025), p. 2580696.

[26] UniProt. UniProt. Jan. 1, 2013. URL: https://www.uniprot.org/proteomes/UP000005640 (visited on 12/21/2025).

[27] Helen M. Berman et al. “The Protein Data Bank”. In: Nucleic Acids Research 28.1 (Jan. 1, 2000), pp. 235–242. ISSN: 0305-1048. DOI: 10.1093/nar/28.1.235. URL: https://doi.org/10.1093/nar/28.1.235 (visited on 12/21/2025).

